# Lotka-Volterra Dynamics Facilitate Sustainable Biocontrol of Wastewater Sludge Bulking

**DOI:** 10.1101/2025.03.29.646071

**Authors:** Fabienne Baltes, Antonia Weiss, Marina Ettl, Kenneth Dumack

## Abstract

Biological wastewater treatment is driven by complex interactions between prokaryotes and eukaryotes. Occasionally, filamentous bacteria species, first and foremost *Ca.* Microthrix parvicella, increase in abundance and lead to detrimental wastewater sludge bulking or floating, causing environmental harm and financial losses. Current mitigation strategies rely heavily on nonspecific chemical interventions, which present environmental risks and lack prolonged effectiveness. Here, we utilise long-term monitoring data from four German wastewater treatment plants (WWTPs) to explore sustainable biocontrol alternatives. Our findings reveal Lotka-Volterra dynamics between *Ca.* Microthrix parvicella and the protist *Arcella* spp. Visual and experimental validation demonstrate the suppression of filamentous bacterial growth by predation. We further model these interactions, predicting the biocontrol potential of *Arcella* spp. for both immediate and sustained efficacy in managing sludge bulking. These results highlight the potential of protists as biological control agents, providing a more sustainable and environmentally friendly alternative to chemical treatment.

## Introduction

Water is a precious and finite resource, and the utilisation of microorganisms in wastewater treatment and reclamation holds immense significance. To recycle wastewater, societies depend on wastewater treatment plants (WWTPs), with a biological treatment step (i.e. the activated sludge basin), where a multitude of microorganisms act in concert ^1–3^. The activated sludge microbiome consists of thousands of species, a core and transient microbiome consisting of bacteria, archaea, protists, fungi, and even microscopic animals. Due to the large scale and partial openness of the bioreactor to the environment this microbiome underlies fluctuations in community composition and functional efficiency. To cope with and steer this microbiome into a desired direction, wastewater treatment plant operators commonly manage the microbiome indirectly through a change in physiochemical parameters, like an adjustment of retention time and aeration, or chemical supplementation ^4,10^.

Functions fulfilled by the wastewater microbiome are manifold. The coupling of denitrification and nitrification reduces wastewater nitrogen levels, while aerobic and anaerobic heterotrophs aid in degrading organic materials ^3,11,12^. The growth of microbial biomass plus aeration leads to flocculation, facilitating the separation of solids through sedimentation in the following clarification step ^2,5,7^. One of the most important challenges in the biological step of wastewater treatment, if not the most critical, is sludge bulking, alongside the related issue of sludge foaming (hereafter referred together to as sludge bulking for simplicity). Both disrupt sedimentation and thereby prevent efficient sludge removal ^10,13–15^. As a result, insufficiently treated wastewater may be released into the environment, which is accompanied by a detrimental environmental impact and, in many countries, leading to heavy fining of wastewater treatment plant operators, if detected ^16,17^. Accordingly, there is a high interest in predicting, treating, and preventing sludge bulking. Common technical control measures are not always successful and most sludge bulking treatments rely on the supplementation of chemical precipitants ^13,14,17^. Additional precipitant usage as control measure is expensive, and the use of non-dedicated agents can negatively impact key functional bacteria, including nitrifiers, thereby impairing the functionality of the WWTP ^14,15,17,18^. It is thus crucial to better understand microbiome functioning in order to develop more sustainable ways of preventing sludge bulking during wastewater treatment.

Sludge bulking is caused by a bloom of filamentous bacteria mostly *Proteobacteria, Actinobacteria, Firmicutes, Planctomycetes, Betaproteobacteria* and *Chloroflexi*, it is recognised when the sludge volume index (SVI) exceeds 150mL/g ^3,15,18^. Globally, the most prominent bacterial genus causing sludge bulking events all over the globe is *Ca.* Microthrix parvicella, belonging to the *Actinobacteria* ^14,19^. To this date, *Ca.* M. parvicella could not be sufficiently cultured, preventing its formal description and the precise measurement of its physiological capabilities and parameters affecting its abundance ^14,20,21^. Nonetheless, it is known that *Ca.* M. parvicella preferentially degrades long-chain fatty acids, grows relatively slowly, and shows its highest abundances in low dissolved oxygen (DO) environments and at colder temperatures (< 20°C) ^14,15,20,22^. However, it is particularly unclear whether the negative correlation of its abundance with temperature stems simply from the physiological constraints of this species‘ metabolism or whether it could be explained by seasonal ecological interactions.

Predation has been observed to impact the microbial community composition and biomass in WWTPs. Predators affect wastewater prokaryotic diversity and taxon richness, evenness, as well as beta diversity (Arregui et al., 2010; Burian et al., 2022; Kim and Unno, 1996; Lee and Welander, 1996a, 1996b). Heck et al. (2023) recently showed that the community composition of wastewater predators, i.e. protists and microscopic animals, was tightly linked to changes in temperature. While wastewater prokaryotes were unaffected by seasonal changes in temperature, their community composition was impacted by the seasonally changing community of temperature-dependent predators. As a result lower variability of the prokaryotic community composition during warm periods was found, most likely due to high predation pressure in wastewater during warmer periods of the year ^27,28^.

In case of *Ca.* M. parvicella, potential biocontrol through predation by rotifers has been intensively investigated. However, despite decades of research, rotiferan biocontrol of *Ca.* M. parvicella never led to full-scale application, presumably due to culturing problems of rotifers, small effect sizes, and/or their sensitivity to toxins and low temperatures ^14,28–30^. Protistan biocontrol potential of *Ca.* M. parvicella remains unexplored since filamentous growth is considered an effective defence mechanism against predation by protists. However recently, Dumack et al. (2024) found evidence that testate amoebae within the Arcellinida (Amoebozoa) evolved special adaptation for the consumption of large and filamentous prey. Arcellinida evolved a shell that acts as an anchor for actin of the pseudopodia to pull on and break large and filamentous prey ^32–34^. Arcellinida are among the most abundant wastewater protists, and although predation on filamentous bacteria was anecdotally reported, their biocontrol potential was never thoroughly assessed ^30,35,36^.

Despite the high needs to prevent the discharge of sludge material into the environment and to reduce the reliance on additional chemical precipitants, no environmentally sustainable biocontrol strategy for sludge bulking is as yet available ^13,14,17^. This study investigates multiple long-term monitoring datasets from municipal German WWTPs to explore the linkage of Arcella spp. and filamentous bacteria in interkingdom wastewater ecology. Special attention was given to filament density, the abundances of filamentous bacteria species, and their correlations with physicochemical and biotic factors. We identified a naturally occurring antagonistic protist to filamentous bacteria and shifted from correlative data exploration to causative, experimental validation and prediction to explore the biocontrol potential of the protist *Arcella* spp..

## Results

Data sampled for almost a decade at a German WWTP revealed that the activated sludge microbiome undergoes regular annual cyclic fluctuations, with a distinct cluster community in the warmer seasons (summer and fall) clearly separated from communities in the colder seasons (winter and spring; Fig. 1A). This pattern indicates a clear change of temperature-dependent recurring annual microbial communities in wastewater. The vector of filament density, a measurement for the abundance of filamentous bacteria, indicated a strong association with colder seasons and the abundance of *Ca.* M. parvicella, showing a strong dependence of filament density on the abundance of *Ca.* M. parvicella with season. A subsequent correlation analysis with significant negative correlations with temperature (R² = - 0.4) between taxa, the filament density, and environmental variables supports the finding that seasonal temperature fluctuations are the main driver of *Ca.* M. parvicella abundance and filament density.

**Figure 1:**
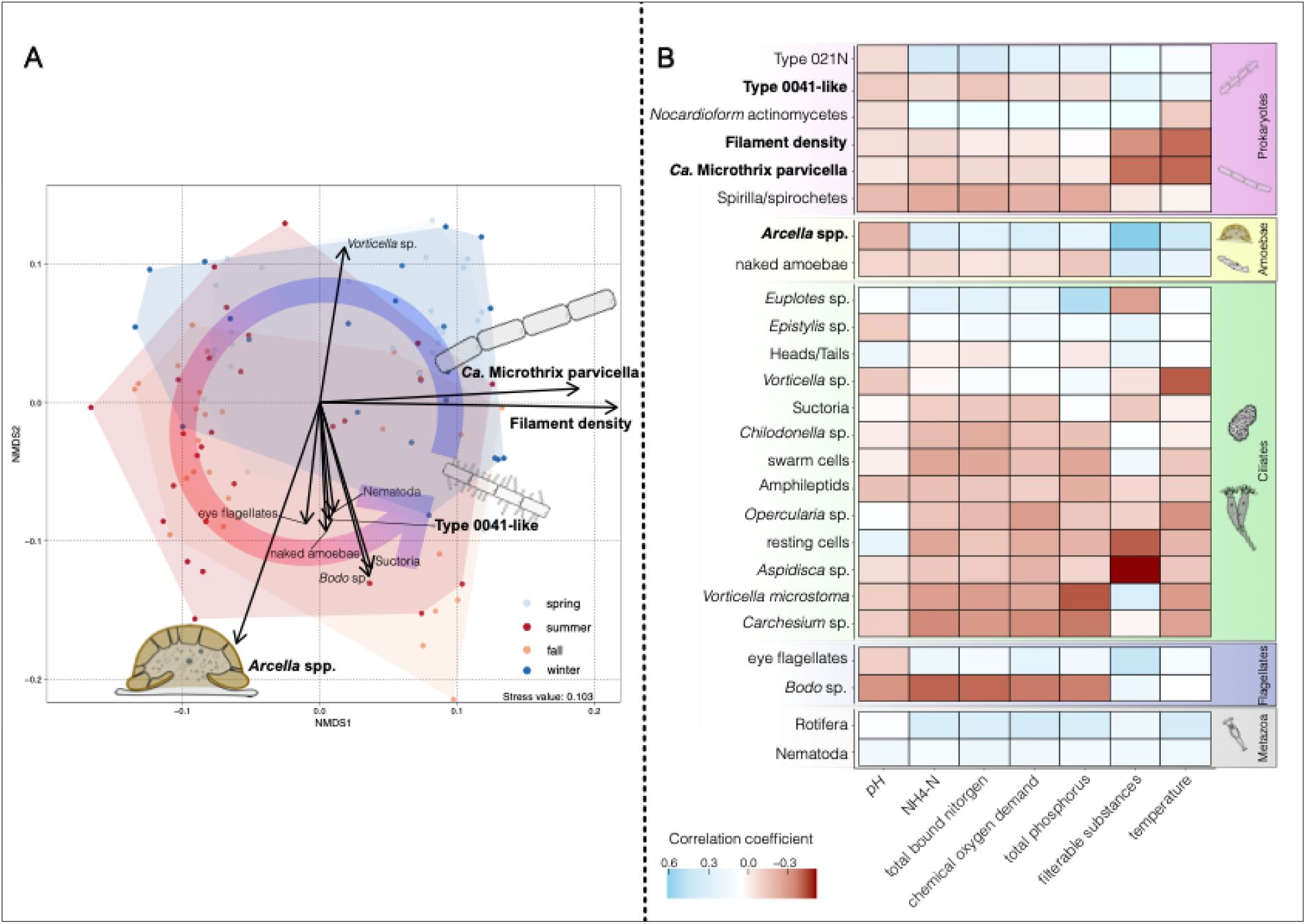
Seasonal dynamics and correlation patterns in the microbial communities of an aerated bioreactor (Dataset 1). (A) Non-metric multidimensional scaling (NMDS) plot, with taxonomic organism variables fitted to the ordination space, illustrating the seasonal dynamics of the microbial community, the round arrow illustrates the annual cycle. The colours of the data points indicate the corresponding season (light blue = spring, blue = winter, red = summer, orange = fall). The length of each arrow represents the strength of the association between the corresponding organism and the ordination space. (B) Heatmap depicting the results of Spearman’s rank correlation analysis between the observed taxa and associated environmental metadata. Blue and red indicate positive and negative correlation coefficients, respectively. Note the strong negative correlation between filament density and the abundance of Ca. M. parvicella and temperature, indicating a temperature dependence of both. Drawings illustrate taxa of particular importance in the subsequent analyses, i.e. Ca. M. parvicella, type 0041-like filamentous bacteria, and Arcella spp..

Notably, while the vectors of most warmer-season-associated taxa pointed in a similar direction, the vector for *Arcella* spp. was distinctly shifted to the opposite direction of the *Ca.* M. parvicella and filament density vectors. This suggests the presence of an additional strong variable, potentially a biotic relationship, influencing the abundance of *Arcella* spp. positively while negatively affecting *Ca.* M. parvicella and filament density.

In order to investigate whether a direct association between both groups existed, we analysed correlations among eukaryotic and prokaryotic taxa (Fig. 2A). Strong negative correlations between *Arcella* spp. and *Ca.* M. parvicella, as well as filament density (correlation coefficients of -0.24 and -0.2, respectively), were revealed. These strong negative correlations, even without considering temperature in the analysis, indicate an inverse relationship between *Arcella* spp. and *Ca.* M. parvicella and filament density.

**Figure 2:**
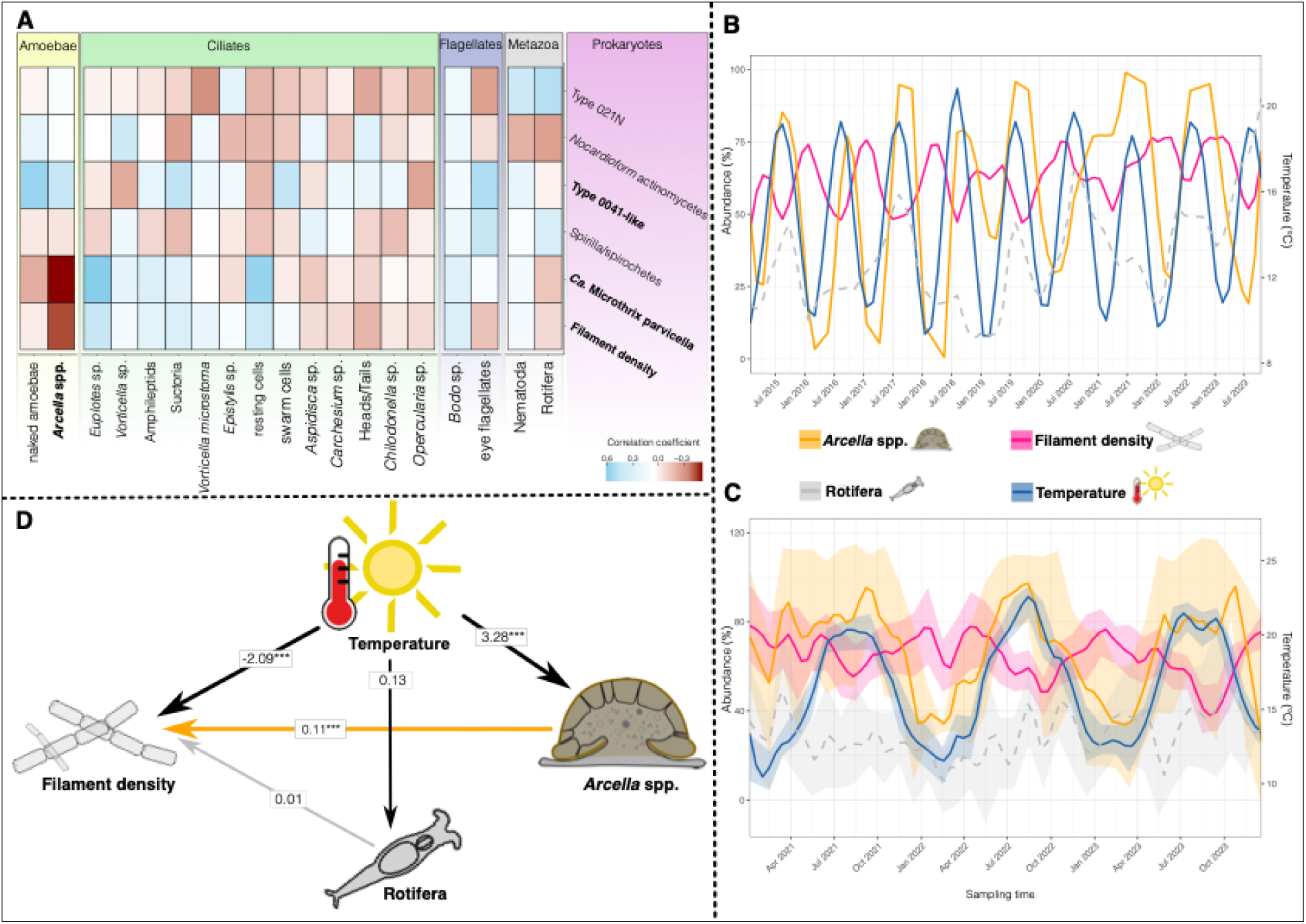
Interactions between Arcella spp. and filamentous bacteria in aerated bioreactors. (A) Heatmap depicting the Spearman’s rank correlation analysis results between eukaryotic and prokaryotic taxa from Dataset 1. Blue and red indicate positive and negative correlation coefficients, respectively. Note the strong negative correlations between Ca. M. parvicella, filament density and Arcella spp. suggesting an inverse relationship between the organisms. (B) Lineplot showing the abundance trends of selected organisms from Dataset 1. The alternating patterns between Arcella spp., temperature, and filament density suggest contrasting interactions, whereas rotifer abundance does not exhibit a clear trend. (C) Lineplot showing abundance patterns in the replicated dataset where variations are less pronounced, but still visible due to noise and variation. The rotifer abundance remains seemingly random. (D) Structural equation modeling (SEM) results demonstrate that temperature acts as the primary driver of changes in filament density and Arcella spp. abundance, while Arcella spp. also exerts an independent influence on filament density. Illustrations are provided for Arcella spp., filament density, and rotifers.

Regular temperature-dependent fluctuations of filament density and the abundance of *Arcella* spp., occurred (Fig. 2B). *Arcella* spp. abundance, as well as filament density, rise and fall with temperature. Notably, filament density appears to peak during periods when *Arcella* spp. abundances are low. Additionally, rotifers as putative antagonists of filamentous wastewater bacteria, were added. Despite a general positive correlation of rotifer abundance with temperature, the correlation of rotifer abundance with filament density was rather weak.

To provide stronger evidence all datasets were aggregated to obtain a replicated time series of spatially independent WWTPs that exhibit the same annual cycling, although showing more variability (Fig. 2C). SEM was used to disentangle the impact of temperature and the abundance of *Arcella* spp. and rotifers on filament density. The abundance of *Arcella* spp. was positively correlated with increasing temperature (path coefficient (pc) = 3.28) and explains a significant proportion of the overall filament density (pc = 0.11) but temperature affected filament density negatively (pc = -2.09). Surprisingly, this effect of *Arcella* spp. on filament density was positive, which may be explained by the nature of time serial data. Note that the path coefficient of 0.11 seems low, however, the abundances of both groups are measured as categorical data meaning that the coefficient in this case must be interpreted in percent, i.e. an increase of about 10 abundance categories of *Arcella* spp. corresponds to an increase of one abundance category of the filament density. Since no significant effect was found for rotifers, they were excluded from further analysis.

The influence of *Arcella* spp. on the filament density was further examined by analysing whether *Arcella* spp. generalistically or selectively affected groups of filamentous bacteria. Only *Ca.* M. parvicella showed strong seasonal fluctuations (Fig 3A), while the abundance of type 0041-like filamentous bacteria was much more stable, suggesting a closer association of *Arcella* spp. with *Ca.* M. parvicella. In support, the SEM reveals a significant impact of *Arcella* spp. on *Ca.* M. parvicella (pc = 0.14), but not on type 0041-like filamentous bacteria. Additionally, type 0041-like filamentous bacteria show a significant negative association with both, filament density (pc = -0.12) and *Ca.* M. parvicella (pc = -0.17), indicating competitive exclusion and that their contribution to filament density was low. In contrast, *Ca.* M. parvicella demonstrates a significant positive association with filament density (pc = 0.48). Furthermore, no direct impact on filament density by *Arcella* spp. was observed, indicating that the effect on filament density (as shown in Fig. 2D) is mediated indirectly through the abundance of *Ca.* M. parvicella. Furthermore, the strong influence of temperature on *Ca.* M. parvicella (pc = - 2.34) supports that the abundance of *Ca*. M. parvicella is the main driver of filament density.

**Figure 3:**
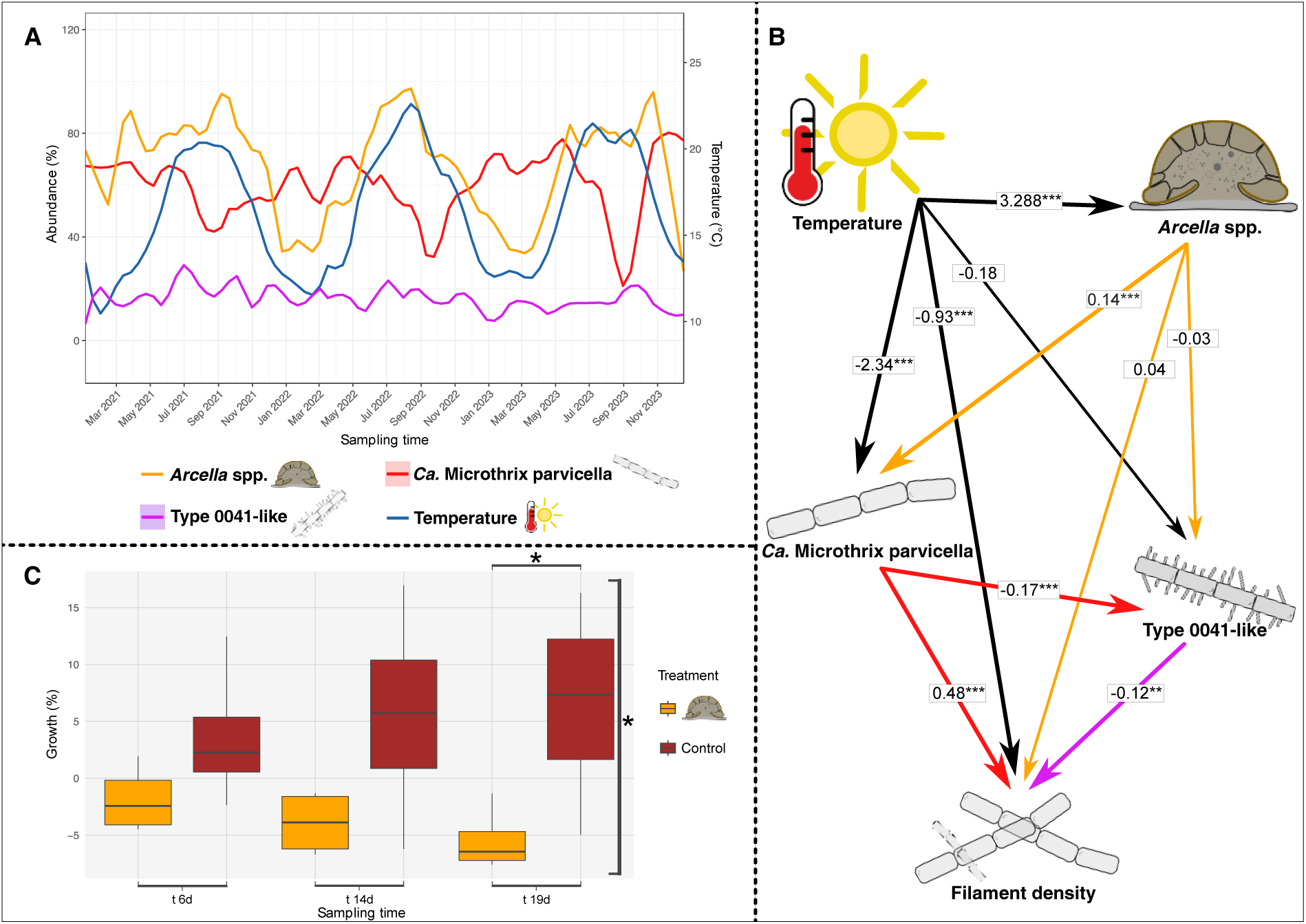
Specific influence of Arcella spp. on filamentous bacteria. (A) Lineplot depicting abundance patterns in the replicated dataset, with Ca. M. parvicella and type 0041-like filamentous bacteria as representatives for filament density. Alternating abundance trends between Arcella spp. and Ca. M. parvicella suggest a potential interaction, while type 0041-like filamentous bacteria show no notable interaction here. (B) SEM results for all considered organisms, highlighting temperature as the primary influencing factor on Arcella spp. and Ca. M. parvicella. A significant negative association is observed between type 0041-like filamentous bacteria and filament density, suggesting a limited role in overall filament formation. In contrast, Ca. M. parvicella shows a significant positive association with filament density, alongside a negative relationship with type 0041-like filamentous bacteria. Additionally, Arcella spp. exhibit a positive association with Ca. M. parvicella abundance, suggesting a specific link between these taxa. (C) Boxplot of the microcosm experiment showing changes in the abundance of filamentous bacteria and floc biomass over time. Arcella treatment samples exhibit significantly reduced filamentous bacterial abundance at the final time point t_19d_, providing strong evidence for Arcella spp. mitigating filamentous bacterial growth. Illustrations are provided for Arcella spp., Ca. M. parvicella, filament density, and type 0041-like filamentous bacteria.

To estimate the type of interaction, its time delay and impact, a microcosm experiment was set up. During 19 days of incubation *Arcella hemisphaerica* not only suppressed bacterial growth (MANOVA; p-value < 0.05), but also reduced the abundance of bacteria and flocs (t-test, p-value < 0.05). This clearly antagonistic predator-prey relationship between the bacterivorous *Arcella* spp. and the filamentous bacterium *Ca.* M. parvicella indicates that *Arcella* spp. serves as a natural biocontrol agent mitigating filamentous bacteria. The predator-prey interaction could be confirmed by video documentation *A. hemisphaerica* consumed up to ten bacterial filaments simultaneously with a handling time of about 30 seconds each (Fig. 4A-F). *A. hemisphaerica* attached to the filament ends with its pseudopodia and subsequently pulled the filaments to the aperture, i.e. the opening of the amoeba’s shell, for phagocytosis. During consumption, some bacterial filaments were bent. The video shows the consumption of one bacterial filament every three seconds by *A. hemisphaerica*. However, these numbers have to be understood as the maximal observed feeding rate observed as feeding in *A. hemisphaerica* is no uniform process. *In situ* observation of activated sludge samples showed *A. hemisphaerica* to ingest several morphologically different bacterial filaments (Fig. 4G-I), including entire attached sludge flocs (Fig. 4H).

**Figure 4:**
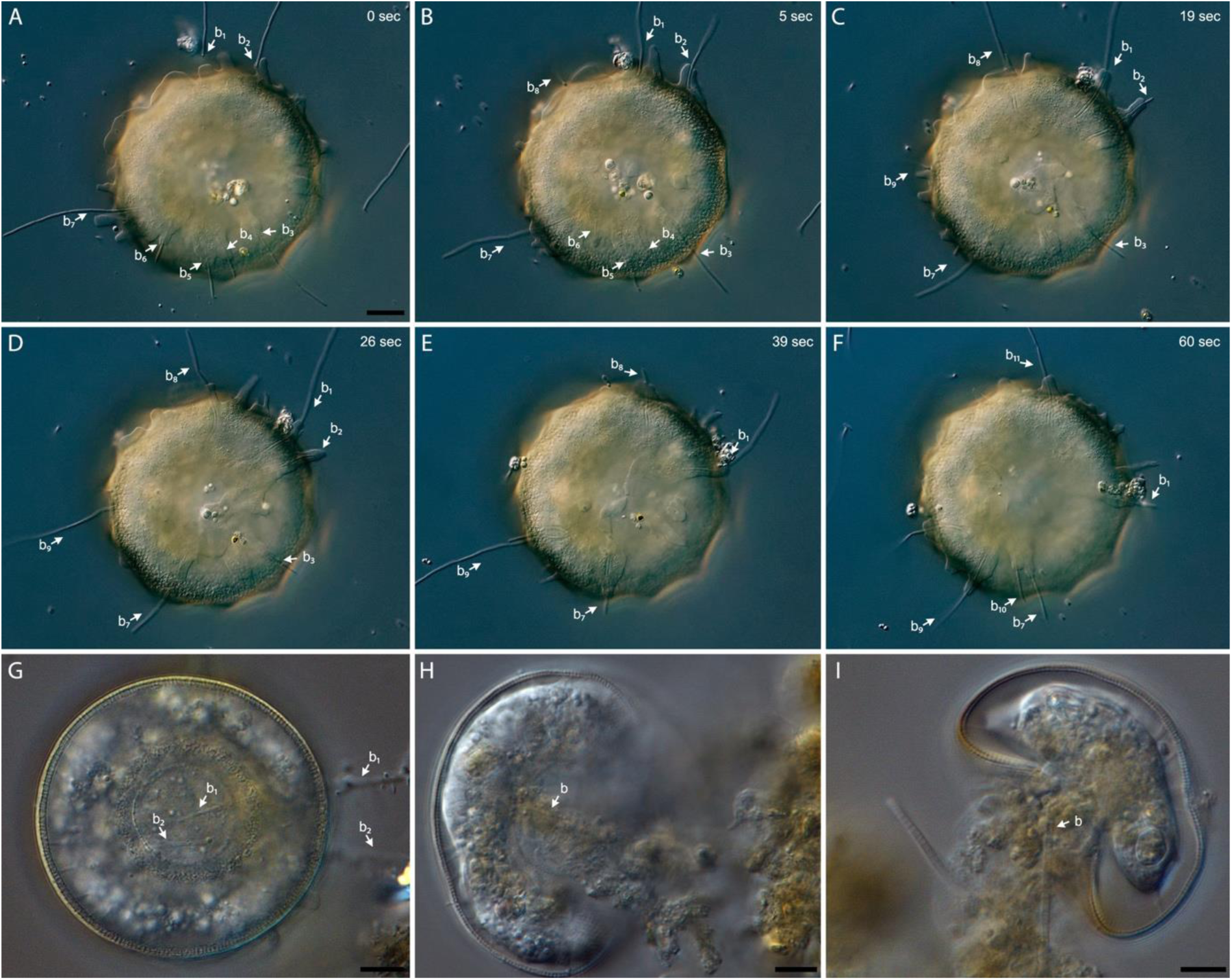
Arcella hemisphaerica feeding on multiple filamentous bacteria and sludge flocs. (A-F) A time series spanning over 60 seconds. Every single bacterial filament is labelled and numbered. (G-I) Morphologically different wastewater filamentous bacteria are ingested. A video is provided as Supplementary Video 1.

**Figure 5:**
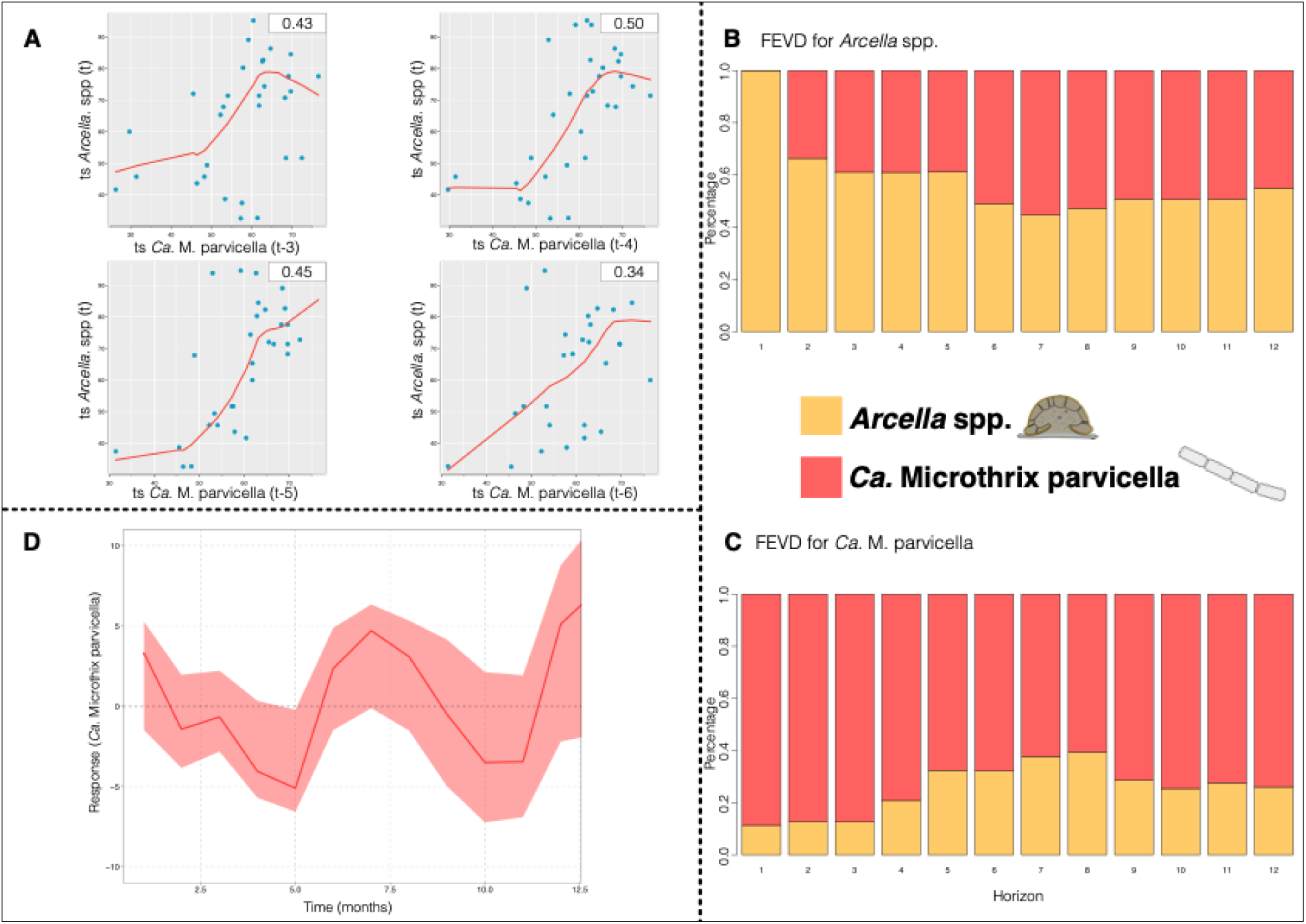
Predator-prey dynamics over time. (A) Lag plots depicting relationships between Ca. M. parvicella (x-variable) and Arcella spp. (y-variable) across different time lags, showing the strongest interactions at time lags t-3, t-4, t-5, and t-6 months. (B, C) FEVD plots for Arcella spp. (B) and Ca. M. parvicella (C), respectively. The variance of prediction errors for both variables is primarily explained by their own previous abundances. The forecast horizon is measured in 1-month intervals along the horizontal axis. (D) IRF showing the response of the Ca. M. parvicella time series to a shock in Arcella spp. abundance. The results indicate a reduction in bacterial abundance over approximately five months. The y-axis quantifies the deviation of the variable from its original equilibrium after the shock.

To disentangle time-dependent correlation and causation, we subjected the data to temporal analyses. Granger causality analysis indicated a directed impact of *Ca.* M. parvicella on *Arcella* spp. (p = 0.047), with a marginally supported (p = 0.09) immediate effect within the same sampling period, i.e. within the same month. Accordingly, we tested for a delayed response to further define the type of association between *Arcella* spp. and *Ca*. M. parvicella. The time lag analysis identified a long-time (annual), temporal relationship between *Ca.* M. parvicella and *Arcella* spp. abundances, showing a positive correlation between the abundances of *Ca.* M. parvicella and *Arcella* spp. at negative time shifts of three to six months (i.e. _t-3_, _t-4_, _t-5_, and _t-6_; Fig. 4A, Fig. S11). These findings suggest that strong changes in *Ca.* M. parvicella abundance precede changes in *Arcella* spp. abundance by approximately three to six months and an immediate impact within one month is weakly supported.

Further, we investigated how changes in the abundance in one of the populations affect the other over time. Forecast error variance decomposition (FEVD) analysis revealed that up to 55.4% of the variance in *Arcella* spp. was explained by the prior abundance of *Ca.* M. parvicella, highlighting a substantial influence of *Ca.* M. parvicella on *Arcella* spp.. Conversely, up to 39.4% of the variance in *Ca. M. parvicella* was attributed to the prior abundance of *Arcella* spp., indicating a notable but somewhat lower explanatory power of *Arcella* spp. on *Ca.* M. parvicella.

It seems that the abundance of *Ca.* M. parvicella has a strong influence on the abundance of *Arcella* spp., which in turn affect *Ca.* M. parvicella to a measurable, but weaker extent. To estimate the biocontrol potential, we subjected the data to an impulse response function (IRF) analysis (Fig. 4C). The model supports a biocontrol potential against an overgrowth of *Ca*. M. parvicella by an addition or increase of *Arcella* spp. abundance through a predicted immediate decline in the abundance of *Ca.* M. parvicella lasting for about five months.

## Discussion

### The Lynx and the Hare

In the early 20th century, biologist Charles Gordon Hewitt observed oscillating, multi-year lagged patterns in the abundances of predatory Canadian lynx and their prey, snowshoe hares, using data derived from fur sales records of the Hudson’s Bay Company. These dynamics followed the dynamics predicted by the Lotka-Volterra equation ^37,38^, and have since become a fundamental example illustrating the principles of predator-prey interactions in ecology. This foundational work has led to the development of more complex models and numerous applications in ecology-based management processes, such as integrated pest management ^39–41^. Based on this background, we were able to leverage large-scale data spanning over multiple years and statistical tools developed for time series ecology to showcase the biocontrol potential of *Arcella* spp. against *Ca.* M. parvicella and sludge bulking. The lynx and hare, whose populations are interconnected demonstrate how predation pressure and resource availability interact to shape population cycles ^42,43^. Similarly, our findings reveal that *Ca.* M. parvicella and *Arcella* spp. exhibit a comparable predator-prey dynamics, mediated by seasonal changes in temperature ^7,44,45^.

Currently, effective strategies to control *Ca.* M. parvicella sludge bulking in WWTPs are limited. Existing approaches, such as reducing sludge retention time (SRT) and controlling DO concentrations ^19,21^, as well as the use of chlorine, polyaluminium chloride products (PAX16 and PAX18), nickel as an heavy metal inhibitor, and synthetic polymers, most come with significant drawbacks ^46–49^. These include unintended impacts on beneficial bacteria, short-lasting effects, and serious environmental side effects ^14–16,50,51^. Ozone treatment and magnetic field applications have also been explored, but require further evaluation ^15,52^. Altogether, these limitations highlight the benefit of an efficient, biological, long-term, and cost-effective solution for controlling *Ca.* M. parvicella.

Our findings suggest that promoting natural predators of *Ca.* M. parvicella could offer an environmentally friendly alternative for managing microbial communities in WWTPs. *Ca.* M. parvicella reveals its highest abundances at colder temperatures, due to reduced predation pressure by *Arcella* spp. that respond sensitive to low temperature ^14,15,20,22^. Time lag analysis suggested that *Ca*. M. parvicella and *Arcella* spp. populations follow the annual cycle tightly as they are lagged by three to six months. The time lag analyses as well as the positive path coefficients in our models demonstrate that changes in the abundance of *Arcella* spp. follow the changes of abundance of *Ca.* M. parvicella under natural conditions. Accordingly, simple increase of SRT to increase Arcella spp abundances is likely to be accompanied by further growth of *Ca*. M. parvicella ^10,21,53^. Instead of a simple increase of SRT, we show that the abundance of *Arcella* spp. must be increased for effective treatment, for instance by the addition of cultured material. An immediate effect of *Ca*. M. parvicella abundances within the same sampling month is supported only weakly with a p-value of less than 0.1. Despite this weak statistical support of an immediate effect within a single month based on environmental data, our experiment confirms that after 19 days subsequent to supplementation, *A. hemisphaerica* did not only lead to a suppression of filament and floc growth, but a significant decline. IRF analyses indicates that after biocontrol application of *Arcella* spp., a long-lasting effect, up to five months, could be achieved. Collectively, these findings suggest an effective biocontrol based on a naturally occurring Lotka-Volterra predator-prey dynamic in the aeration tanks of WWTPs by supplementation.

Our findings extend previous work on rotifers as biocontrol agents ^29,54^. In contrast to rotifers, *Arcella* spp. demonstrate consistent, long-term associations with *Ca.* M. parvicella populations across full-scale WWTPs, suggesting greater potential for practical application. Now, the next step to this research is a proof of concept of biocontrol on a full-scale WWTP with industrial-scale addition of *Arcella* spp. or engineering facilities to specifically increase naturally occurring *Arcella* spp. abundances. While doing so, we recommend to investigate potential side effects caused by a high grazing pressure of *Arcella* spp. on sludge composition, abundance and density of flocs, and subsequent affection of sedimentation behaviour.

## Conclusion

Promoting *Arcella* spp. populations in treatment systems could provide a sustainable solution to manage sludge bulking and improve treatment efficiency without harmful chemical additives. Future research should focus on full-scale trials to validate *Arcella* spp.-based biocontrol in WWTPs and explore optimal conditions for *Arcella* spp. proliferation. By integrating these biological control strategies, WWTPs could reduce sludge bulking, leading to more resilient and efficient wastewater treatment processes and cost efficiency.

## Material and Methods

### Data Collection and Preparation

Data were accessed from four WWTPs across Germany, referred to as Datasets 1, 2, 3, and 4 to maintain requested anonymity. Staff trained in morphological identification of wastewater bioindicators gathered the data through regular microscopic examination of samples obtained from aeration tanks, according to the guidelines outlined by the Bavarian Environment Agency ^10^. These guidelines recommend the quantification of the abundance of prokaryotic and eukaryotic microorganisms with bioindication capacity as well as the filament density as a rapid assessment of WWTP functionality. The abundance data provided by each of the facilities were reported as categorical values. For testate amoebae, for example, category 0 represents 0 individuals, category 1 represents 1-5 individuals, category 2 represents 6-10 individuals, and category 3 represents 11 or more individuals per sample. Environmental parameters were considered in our analysis when available, with water temperature being measured consistently across all monitoring datasets. Other parameters varied by dataset: Dataset 1 included pH value, ammonium, chemical oxygen demand (COD), total phosphorus concentration, total bound nitrogen concentration (TNb), total organic carbon (TOC), and filterable substances; Dataset 2 only included pH value, nitrite, nitrate, ammonium, total nitrogen, and TNb; and Datasets 3 and 4 included the sludge volume index (SVI). Sampling frequency and duration also differed across datasets, with Dataset 3 being sampled weekly for 35 months, while Datasets 1, 2, and 4 were sampled monthly, spanning 102, 66, and 35 months, respectively (Supplementary Table S1).

All statistical analyses were conducted using R version 4.3.1 ^55^ in RStudio, employing the packages vegan v. 2.6.4 ^56^, tidyverse v. 2.0.0 ^57^, dplyr v. 1.1.3 ^58^, lubridate v. 1.9.3 ^59^, readxl v. 1.4.3 ^60^, and zoo v. 1.8.12 ^61^. Unless stated otherwise, these packages were used for all data processing and statistical analyses. All scripts and raw data for the subsequently detailed analyses are available on GitHub (Lotka-Volterra-Dynamics-Facilitate-Sustainable-Biocontrol-of-Wastewater-Sludge-Bulking).

### Non-Metric Multidimensional Scaling

Community data were plotted using metaMDS and envfit functions in the vegan package to visualise the strongest drivers of environmental factors and species. Only the top ten variables with the highest explained variation (r² values) and p-values below 0.05 were displayed. Given the categorical nature of the measured abundance data, Gower’s distance was selected as the dissimilarity measure. This metric is particularly well-suited for mixed-type data, as it accommodates both continuous and categorical variables, allowing for an effective quantification of differences between samples ^62^.

### Correlation Analysis and Heatmap Generation

Relationships between environmental parameters and microbial abundances were analysed using Spearman’s rank correlation ^63^. Initial visual inspection of histograms suggested non-normal distributions, which were confirmed through the Shapiro-Wilk test ^64^. Correlations were computed using the cor.test function in R, and the resulting associations were visualised as heatmaps generated with ggplot2 v. 3.4.3 ^65^ and reshape2 v. 1.4.4 ^66^.

### Assessment of Temporal Fluctuations

Lineplots were used to visually assess temporal fluctuations in the long-term Dataset 1 and the replicated dataset. The replicated dataset was obtained by the aggregation of Datasets 1-4 with at least three samples per month covering 2021 to 2023 (n = 249), which allowed for a comparison of monthly sampling intervals. Only selected organisms, *Ca.* M. parvicella, “type 0041-like filamentous bacteria”, *Arcella* spp., rotifers, the “filament density,” and temperature were included in the replicated dataset. In this context, “filament density” refers to the sum of all individual filaments observed in the sample (German: “Gesamtfädigkeit”) ^10^, while type 0041-like filamentous bacteria represent bacterial filaments of unknown taxonomy, colloquially called “type 0041” and similar morphotypes of unknown taxa ^13,17^.

### Structural Equation Modeling

Structural equation modeling (SEM) was used to test and refine hypothesised models of ecosystem processes and interactions ^67^. Models were calculated using the lavaan package v.0.6.18 ^68^, and visualised using tidySEM v. 0.2.7 ^69^.

### Mechanistic Background Evaluation and Laboratory Experiment

In addition, direct microscopic investigations to confirm the interaction of *Arcella* spp. with filamentous bacteria were implied. For this, *Arcella hemisphaerica* was observed in natural samples from (a) a pond (coordinates 52.1253080, 21.0452141) and (b) the aerated sludge bioreactor of the WWTP in Cologne Weiden, Germany (coordinates 50.9391141, 6.8113971). The latter was enriched with a previously established *Arcella hemisphaerica* culture derived from the same WWTP to increase cell numbers. Samples were investigated and continuously filmed with a Nikon Eclipse 90i microscope at 600x and differential interference contrast (Fig. 4).

A microcosm experiment was conducted to experimentally confirm and quantify the putative antagonistic effect of *Arcella* spp. on the abundance of filamentous bacteria. Eight wells of a 24-well-plate were inoculated with 1 ml of wastewater and 20 µl of freshly sampled sludge flocs containing filamentous bacteria. The sludge was sampled from the same WWTP in Cologne Weiden, Germany, as described above on October 28th 2024, and incubated at 15°C one day prior to the experimental setup. Approximately 30 individuals (200μl) of an *A. hemisphaerica* culture derived from the same WWTP, were added as predator treatment (n=4) and 200µl of supernatant culture medium were added to the control (n=4). The microcosms were allowed to sediment for one day before initial documentation (called time point 0 days, i.e. *t_0_*). Changes in optical density (OD) were documented after 6, 14, and 19 days with an inverted microscope and 4x magnification respectively. The pictures were analysed for changes in OD (%) with ImageJ ^70^. OD was determined by a subtraction of brown pixels and an area measurement. Differences in OD with respect to *t_0_* were calculated. Multivariate Analysis of Variance (MANOVA) and t-tests ^71,72^ of respective time points were calculated to evaluate changes in OD due to predation.

### Time Series Analysis

To reduce sampling errors, for the time series analysis, the mean of at least three sampling points from the replicated dataset was calculated to represent each monthly value, resulting in an overall sample size of n = 36.

### Time Lag Analysis

Time lag analysis was employed to disentangle temporal dependencies between *Arcella* spp. and *Ca.* M. parvicella. The lag2.plot function from the astsa package v. 2.1 ^73^ was used to visualise the relationship between these variables at different time lags. This analysis calculated correlations between the y-variable (*Arcella* spp.) at time t and the x-variable (*Ca.* M. parvicella) at prior time points, identifying the optimal lag (in months) based on the highest observed correlation values (Fig. 4A, Fig. S11).

### Vector Autoregression

Vector autoregression (VAR) models were constructed from the time series objects to quantify the reciprocal impact of taxa. The R packages tseries v. 0.10.57 ^74^, vars v. 1.6.1 ^75^, forecast v.8.23.0 ^76^, urca v. 1.3.4 ^77^, and mFilter v. 0.1.5 ^78^ were used to perform VAR modeling and data visualisation. Granger causality tests were conducted, using the causality function (package vars) to evaluate whether past densities of *Ca*. M. parvicella could predict *Arcella* spp. abundance, and *vice versa*. Instantaneous causality was also assessed with this function.

To evaluate species-specific responses to unexpected changes in the other species’ abundance, impulse response function (IRF) analysis was performed using the irf function (package vars)(Fig. 4C)^40^. Additionally, forecast error variance decomposition (FEVD) was employed with the fevd function (package vars), to quantify how much of the variance in prediction errors for one organism could be attributed to changes in the other species’ abundance (Fig. 4B)^79^.

## Author contributions

F.B. and K.D. conceived and designed the study, performed the data analysis, and wrote the manuscript in consultation with M.E.

K.D. supervised the project and contributed to the interpretation of the data.

M.E. contributed to the methodology and provided expertise in the data collection process.

A.W. performed the experimental work and conducted the photography.

All authors read, commented and approved the manuscript.

## Competing interests

The authors declare no competing interests.

## Data availability statement

The raw data are available in the Lotka-Volterra Dynamics Facilitate Sustainable Biocontrol of Wastewater Sludge Bulking repository on GitHub [https://github.com/fabib1209/Lotka-Volterra-Dynamics-Facilitate-Sustainable-Biocontrol-of-Wastewater-Sludge-Bulking/tree/main]. Supplementary materials can be found in the supplementary section of this article.

## Code availability statement

The raw code is also available in the Lotka-Volterra Dynamics Facilitate Sustainable Biocontrol of Wastewater Sludge Bulking repository on GitHub [https://github.com/fabib1209/Lotka-Volterra-Dynamics-Facilitate-Sustainable-Biocontrol-of-Wastewater-Sludge-Bulking/tree/main].

## Acknowledgements

We are indebted to the diverse, anonymous wastewater treatment plant operators who provided the generated data and James Weiss for supporting us with microscopical images.

## Notes

### Competing Interest Statement

The authors have declared no competing interest.

## References

1. Aragaw, T. A. Functions of various bacteria for specific pollutants degradation and their application in wastewater treatment: a review. Int. J. Environ. Sci. Technol. 18, 2063–2076 (2021).

2. Arregui, L., Pérez-Uz, B., Salvadó, H. & Serrano, S. Progresses on the Knowledge about the Ecological Function and Structure of the Protists Community in Activated Sludge Wastewater Treatment Plants. in vol. 2 972–979 (2010).

3. Ferrera, I. & Sánchez, O. Insights into microbial diversity in wastewater treatment systems: How far have we come? Biotechnology Advances 34, 790–802 (2016).

4. Sobczyk, M., Pajdak-Stós, A., Fiałkowska, E., Sobczyk, Ł. & Fyda, J. Multivariate analysis of activated sludge community in full-scale wastewater treatment plants. Environ Sci Pollut Res 28, 3579–3589 (2021).

5. Freudenthal, J., Ju, F., Bürgmann, H. & Dumack, K. Microeukaryotic gut parasites in wastewater treatment plants: diversity, activity, and removal. Microbiome 10, 27 (2022).

6. Sasi, R. & Suchithra, T. V. Wastewater microbial diversity versus molecular analysis at a glance: a mini-review. Braz J Microbiol 54, 3033–3039 (2023).

7. Heck, N., Freudenthal, J. & Dumack, K. Microeukaryotic predators shape the wastewater microbiome. Water Research 242, 120293 (2023).

8. Wu, L. et al. Global diversity and biogeography of bacterial communities in wastewater treatment plants. Nat Microbiol 4, 1183–1195 (2019).

9. Sun, C. et al. Seasonal dynamics of the microbial community in two full-scale wastewater treatment plants: Diversity, composition, phylogenetic group based assembly and co-occurrence pattern. Water Research 200, 117295 (2021).

10. Pinther, W. et al. Das mikroskopische Bild bei der biologischen Abwasserreinigung. (Bayerisches Landesamt für Umwelt (LfU), 2022).

11. Petrovski, S., Rice, D. T. F., Batinovic, S., Nittami, T. & Seviour, R. J. The community compositions of three nitrogen removal wastewater treatment plants of different configurations in Victoria, Australia, over a 12-month operational period. Appl Microbiol Biotechnol 104, 9839–9852 (2020).

12. Rodríguez, E., García-Encina, P. A., Stams, A. J. M., Maphosa, F. & Sousa, D. Z. Meta-omics approaches to understand and improve wastewater treatment systems. Rev Environ Sci Biotechnol 14, 385–406 (2015).

13. Deepnarain, N. et al. Artificial intelligence and multivariate statistics for comprehensive assessment of filamentous bacteria in wastewater treatment plants experiencing sludge bulking. Environmental Technology & Innovation 19, 100853 (2020).

14. Fan, N.-S., Qi, R., Huang, B.-C., Jin, R.-C. & Yang, M. Factors influencing Candidatus Microthrix parvicella growth and specific filamentous bulking control: A review. Chemosphere 244, 125371 (2020).

15. Syamimi Zaidi, N., et al. Insights into the potential application of magnetic field in controlling sludge bulking and foaming: A review. Bioresource Technology 358, 127416 (2022).

16. Bahrodin, M. B. et al. Recent Advances on Coagulation-Based Treatment of Wastewater: Transition from Chemical to Natural Coagulant. Curr Pollution Rep 7, 379–391 (2021).

17. Sam, T., Le Roes-Hill, M., Hoosain, N. & Welz, P. J. Strategies for Controlling Filamentous Bulking in Activated Sludge Wastewater Treatment Plants: The Old and the New. Water 14, 3223 (2022).

18. Guo, F. & Zhang, T. Profiling bulking and foaming bacteria in activated sludge by high throughput sequencing. Water Research 46, 2772–2782 (2012).

19. Zhang, M., Yao, J., Wang, X., Hong, Y. & Chen, Y. The microbial community in filamentous bulking sludge with the ultra-low sludge loading and long sludge retention time in oxidation ditch. Sci Rep 9, 13693 (2019).

20. Fan, N. et al. Factors affecting the growth of *Microthrix parvicella*: Batch tests using bulking sludge as seed sludge. Science of The Total Environment 609, 1192–1199 (2017).

21. Rossetti, S., Tomei, M. C., Nielsen, P. H. & Tandoi, V. “ *Microthrix parvicella* ”, a filamentous bacterium causing bulking and foaming in activated sludge systems: a review of current knowledge. FEMS Microbiol Rev 29, 49–64 (2005).

22. Wang, J., Li, Q., Qi, R., Tandoi, V. & Yang, M. Sludge bulking impact on relevant bacterial populations in a full-scale municipal wastewater treatment plant. Process Biochemistry 49, 2258–2265 (2014).

23. Burian, A. et al. Predation increases multiple components of microbial diversity in activated sludge communities. The ISME Journal 16, 1086–1094 (2022).

24. Kim, T.-D. & Unno, H. The roles of microbes in the removal and inactivation of viruses in a biological wastewater treatment system. Water Science and Technology 33, 243–250 (1996).

25. Lee, N. M. & Welander, T. Reducing sludge production in aerobic wastewater treatment through manipulation of the ecosystem. Water Research 30, 1781–1790 (1996).

26. Lee, N. M. & Welander, T. Use of protozoa and metazoa for decreasing sludge production in aerobic wastewater treatment. Biotechnology Letters 18, 429–434 (1996).

27. Heck, N., Freudenthal, J. & Dumack, K. Microeukaryotic predators shape the wastewater microbiome. Water Research 242, 120293 (2023).

28. Kocerba-Soroka, W. et al. Lecane tenuiseta rotifers improves activated sludge settleability in laboratory scale SBR system at 13°C and 20°C. Water and Environment Journal 31, 113–119 (2017).

29. Pajdak-Stós, A., Kocerba-Soroka, W., Fyda, J., Sobczyk, M. & Fiałkowska, E. Foam-forming bacteria in activated sludge effectively reduced by rotifers in laboratory- and real-scale wastewater treatment plant experiments. Environ Sci Pollut Res 24, 13004–13011 (2017).

30. Fiałkowska, E. & Pajdak-Stós, A. The role of *Lecane* rotifers in activated sludge bulking control. Water Research 42, 2483–2490 (2008).

31. Dumack, K. et al. It’s time to consider the Arcellinida shell as a weapon. European Journal of Protistology 92, 126051 (2024).

32. Dumack, K., Bonkowski, M., Clauß, S. & Völker, E. Phylogeny and redescription of the testate amoeba Diaphoropodon archeri (Chlamydophryidae, Thecofilosea, Cercozoa), De Saedeleer 1934, and annotations on the polyphyly of testate amoebae with agglutinated tests in the Cercozoa. Journal of Eukaryotic Microbiology 65, 308–314 (2018).

33. Estermann, A. H. et al. Fungivorous protists in the rhizosphere of Arabidopsis thaliana – Diversity, functions, and publicly available cultures for experimental exploration. Soil Biology and Biochemistry 187, e109206 (2023).

34. Siemensma, F. & Opitz, A. M. Beobachtungen an Pseudonebela africana, einer seltenen, doch weltweit verbreiteten Schalenamoebe. Mikrokosmos 103, 248–255 (2014).

35. Hu, B., Qi, R. & Yang, M. Systematic analysis of microfauna indicator values for treatment performance in a full-scale municipal wastewater treatment plant. Journal of Environmental Sciences 25, 1379–1385 (2013).

36. Madoni, P. Protozoa in wastewater treatment processes: A minireview. Italian Journal of Zoology 78, 3–11 (2011).

37. Lotka, A. J. Elements of Physical Biology. (Williams & Wilkins Company, Baltimore, 1925).

38. Volterra, V. Variazioni e fluttuazioni del numero d’individui in specie animali conviventi. Memorie della R. Accademia dei Lincei 6, 31–113 (1926).

39. Deguine, J.-P. et al. Integrated pest management: good intentions, hard realities. A review. Agron. Sustain. Dev. 41, 38 (2021).

40. Ewing, B. T., Riggs, K. & Ewing, K. L. Time series analysis of a predator–prey system: Application of VAR and generalized impulse response function. Ecological Economics 60, 605–612 (2007).

41. Uechi, H., Uechi, L. & Uechi, S. T. The Lynx and Hare Data of 200 Years as the Nonlinear Conserving Interaction Based on Noether’s Conservation Laws and Stability. Journal of Applied Mathematics and Physics 9, 2807–2847 (2021).

42. Krebs, C. J. et al. Impact of Food and Predation on the Snowshoe Hare Cycle. Science 269, 1112–1115 (1995).

43. Stenseth, N. Chr., Falck, W., Bjørnstad, O. N. & Krebs, C. J. Population regulation in snowshoe hare and Canadian lynx: Asymmetric food web configurations between hare and lynx. Proc. Natl. Acad. Sci. U.S.A. 94, 5147–5152 (1997).

44. Flowers, J. J., Cadkin, T. A. & McMahon, K. D. Seasonal bacterial community dynamics in a full-scale enhanced biological phosphorus removal plant. Water Research 47, 7019–7031 (2013).

45. Peces, M. et al. Microbial communities across activated sludge plants show recurring species-level seasonal patterns. ISME Communications 2, 18 (2022).

46. Juang, D. Effects of synthetic polymer on the filamentous bacteria in activated sludge. Bioresource Technology 96, 31–40 (2005).

47. Miłobędzka, A. & Muszyński, A. Selection of methods for activated sludge bulking control using a molecular biology technique combined with respirometric tests. bta 3, 187–193 (2016).

48. Nielsen, P. H. et al. Control of Microthrix parvicella in Activated Sludge Plants by Dosage of Polyaluminium Salts: Possible Mechanisms. Acta hydrochimica et hydrobiologica 33, 255–261 (2005).

49. Wang, L. et al. Effects of Ni2+ on the characteristics of bulking activated sludge. Journal of Hazardous Materials 181, 460–467 (2010).

50. Durban, N., Juzan, L., Krier, J. & Gillot, S. Control of Microthrix parvicella by aluminium salts addition. Water Science and Technology 73, 414–422 (2015).

51. Roels, T., Dauwe, F., Van Damme, S., De Wilde, K. & Roelandt, F. The influence of PAX-14 on activated sludge systems and in particular on Microthrix parvicella. Water Science and Technology 46, 487–490 (2002).

52. Levén, L., Wijnbladh, E., Tuvesson, M., Kragelund, C. & Hallin, S. Control of Microthrix parvicella and sludge bulking by ozone in a full-scale WWTP. Water Sci Technol 73, 866– 872 (2016).

53. Lemmer, H., Griebe, T. & Flemming, H.-C. *Ökologie der Abwasserorganismen*. (Springer Berlin Heidelberg, Berlin, Heidelberg, 1996). doi:10.1007/978-3-642-61423-1.

54. Pajdak-Stós, A. & Fiałkowska, E. The Influence of Temperature on the Effectiveness of Filamentous Bacteria Removal from Activated Sludge by Rotifers. Water Environment Research 84, 619–625 (2012).

55. R Core Team. R: A Language and Environment for Statistical Computing. (R Foundation for Statistical Computing, Vienna, Austria, 2023).

56. Oksanen, J., et al. Vegan: Community Ecology Package. (2022).

57. Wickham, H. et al. Welcome to the tidyverse. Journal of Open Source Software 4, 1686 (2019).

58. Wickham, H., François, R., Henry, L., Müller, K. & Vaughan, D. Dplyr: A Grammar of Data Manipulation. (2023).

59. Grolemund, G. & Wickham, H. Dates and Times Made Easy with lubridate. Journal of Statistical Software 40, 1–25 (2011).

60. Wickham, H. & Bryan, J. Readxl: Read Excel Files. (2023).

61. Zeileis, A. & Grothendieck, G. zoo: S3 Infrastructure for Regular and Irregular Time Series. Journal of Statistical Software 14, 1–27 (2005).

62. Gower, J. C. A General Coefficient of Similarity and Some of Its Properties. Biometrics 27, 857 (1971).

63. Best, D. J. & Roberts, D. E. Algorithm AS 89: The Upper Tail Probabilities of Spearman’s Rho. Journal of the Royal Statistical Society. Series C (Applied Statistics) 24, 377–379 (1975).

64. Royston, J. P. Algorithm AS 181: The W Test for Normality. Applied Statistics 31, 176 (1982).

65. Wickham, H. *Ggplot2: Elegant Graphics for Data Analysis*. (Springer International Publishing : Imprint: Springer, Cham, 2016). doi:10.1007/978-3-319-24277-4.

66. Wickham, H. Reshaping Data with the reshape Package. Journal of Statistical Software 21, 1–20 (2007).

67. Arhonditsis, G. B. et al. Exploring ecological patterns with structural equation modeling and Bayesian analysis. Ecological Modelling 192, 385–409 (2006).

68. Rosseel, Y. lavaan: An R Package for Structural Equation Modeling. Journal of Statistical Software 48, 1–36 (2012).

69. Lissa, C. J. van. tidySEM: Tidy Structural Equation Modeling. (2024).

70. Schneider, C. A., Rasband, W. S. & Eliceiri, K. W. NIH Image to ImageJ: 25 years of image analysis. Nature Methods 9, 671–675 (2012).

71. Legendre, P. & Legendre, L. Numerical Ecology. (Elsevier, 2012).

72. Quinn, G. P. & Keough, M. J. Experimental Design and Data Analysis for Biologists. (Cambridge University Press, 2002).

73. Stoffer, D. & Poison, N. astsa: Applied Statistical Time Series Analysis. (2024).

74. Trapletti, A. & Hornik, K. Tseries: Time Series Analysis and Computational Finance. (2024).

75. Pfaff, B. VAR, SVAR and SVEC Models: Implementation Within R Package vars. Journal of Statistical Software 27, (2008).

76. Hyndman, R., et al. forecast: Forecasting Functions for Time Series and Linear Models. 8.23.0 10.32614/CRAN.package.forecast (2009).

77. Pfaff, B. Analysis of Integrated and Cointegrated Time Series with R. (Springer, New York, 2008).

78. Balcilar, M. mFilter: Miscellaneous Time Series Filters. (2019).

79. Lütkepohl, H. New Introduction to Multiple Time Series Analysis: With … 36 Tables. (Springer, Berlin Heidelberg, 2007).

