## Supplementary Material for "Lotka-Volterra Dynamics Facilitate Sustainable Biocontrol of Wastewater Sludge Bulking"

***Table S1:*** *Information on the datasets from 4 different German WWTPs.*

| ***Dataset name*** | **Samples** | **Frequency** | **Period** |
|---|---|---|---|
| *Dataset 1* | 103 | Monthly | 01.2015-10.2023 |
| *Dataset 2* | 72 | Monthly | 01.2018-09.2023 |
| *Dataset 3* | 153 | Weekly | 01.2021-12.2023 |
| *Dataset 4* | 36 | Monthly | 01.2021-12.2023 |

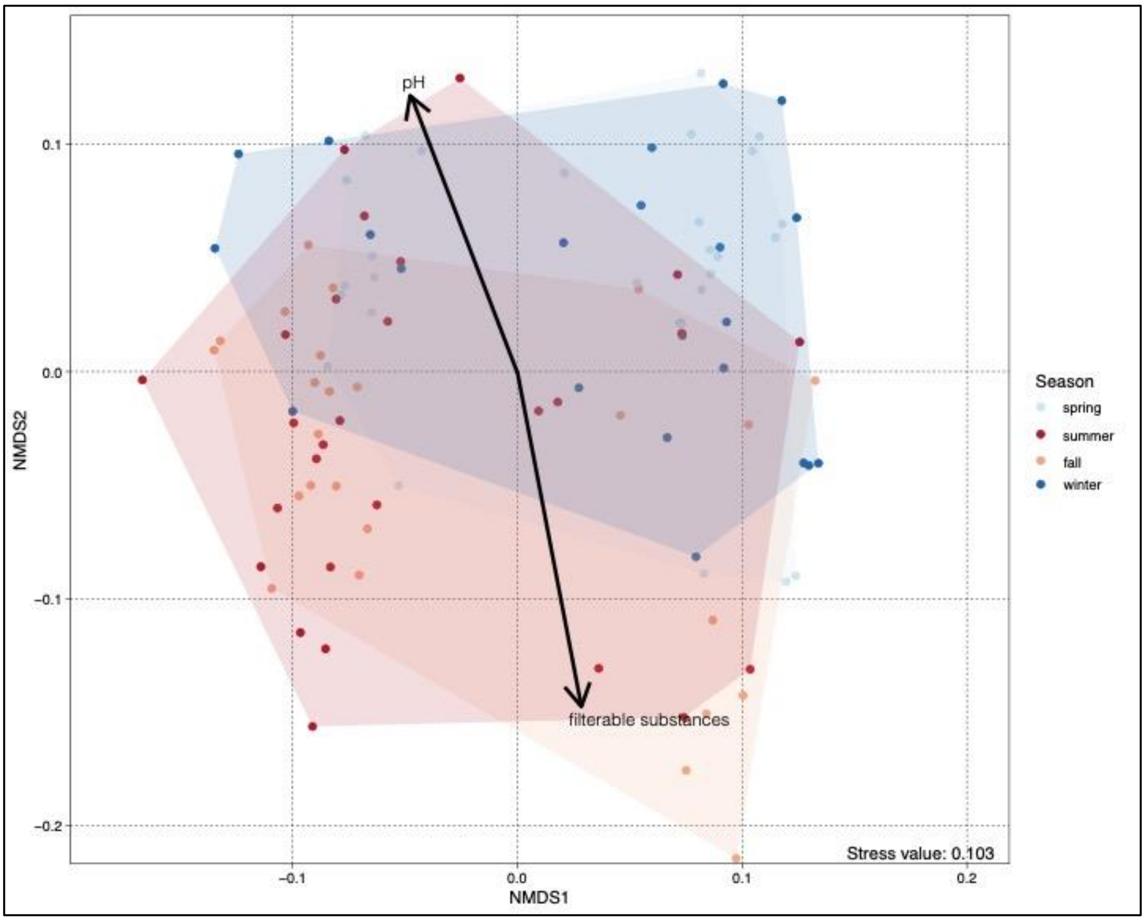


***Figure S1: Abiotic influences in the microbial communities of an aerated bioreactor (Dataset 1)****. Non-metric multidimensional scaling (NMDS) plot, with metadata variables fitted to the ordination space. The colours of the data points indicate the corresponding season (light blue = spring, blue = winter, red = summer, orange = fall). The length of each arrow represents the strength of the association between the corresponding metadata variable and the ordination space. Filterable*

*substances show a strong correlation with the warmer seasons (summer/fall) and pH shows a strong correlation with the colder seasons (winter/spring).*

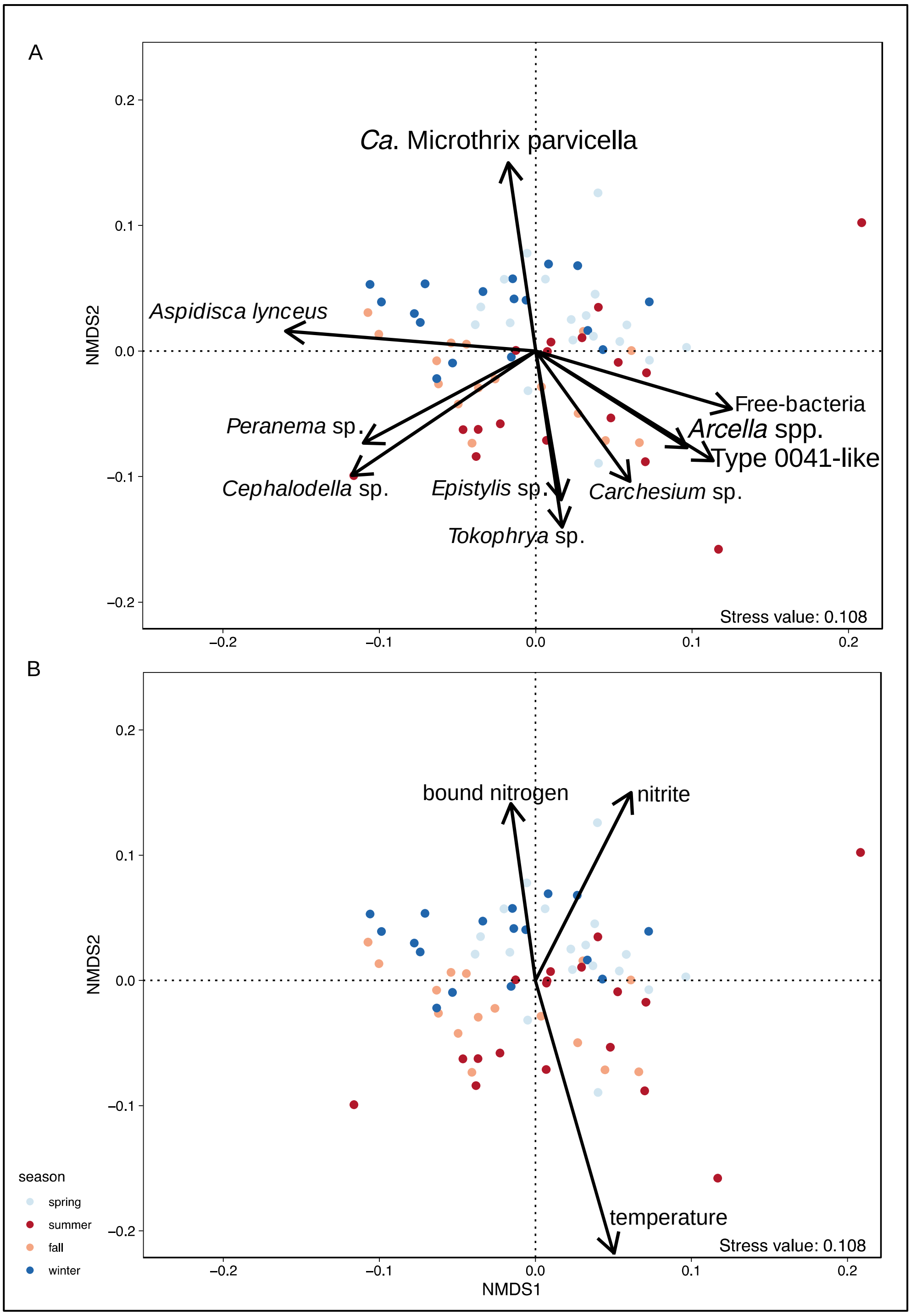


***Figure S2: Seasonal dynamics in the microbial communities of an aerated bioreactor (Dataset 2)****. (A) Non-metric multidimensional scaling (NMDS) plot, with taxonomic organism variables fitted to the ordination space, illustrating the seasonal dynamics of the microbial community. The colours of the data points indicate the corresponding season (light blue = spring, blue = winter, red = summer, orange = fall). The length of each arrow represents the strength of the association between the corresponding organism and the ordination space. (B) Non-metric multidimensional scaling (NMDS) plot, with metadata variables fitted to the ordination space. Ca. M. parvicella shows a strong negative association with temperature and a positive*

association with bound nitrogen and nitrite. Arcella spp. and type 0041-like filamentous bacteria show a more positive association with temperature.*

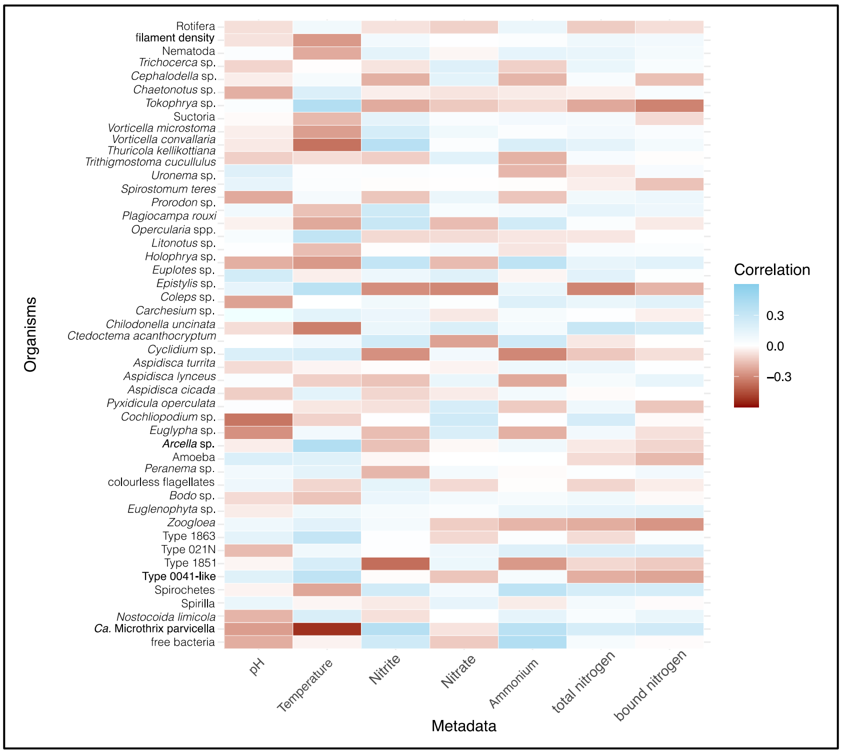


***Figure S3:*** *Heatmap depicting the results of a Spearman's rank correlation analysis for Dataset 2 between the observed taxa and associated environmental metadata. Blue and red indicate positive and negative correlation coefficients, respectively. Note the strong negative correlation between the abundance of Ca. M. parvicella and temperature, indicating a temperature dependence.*

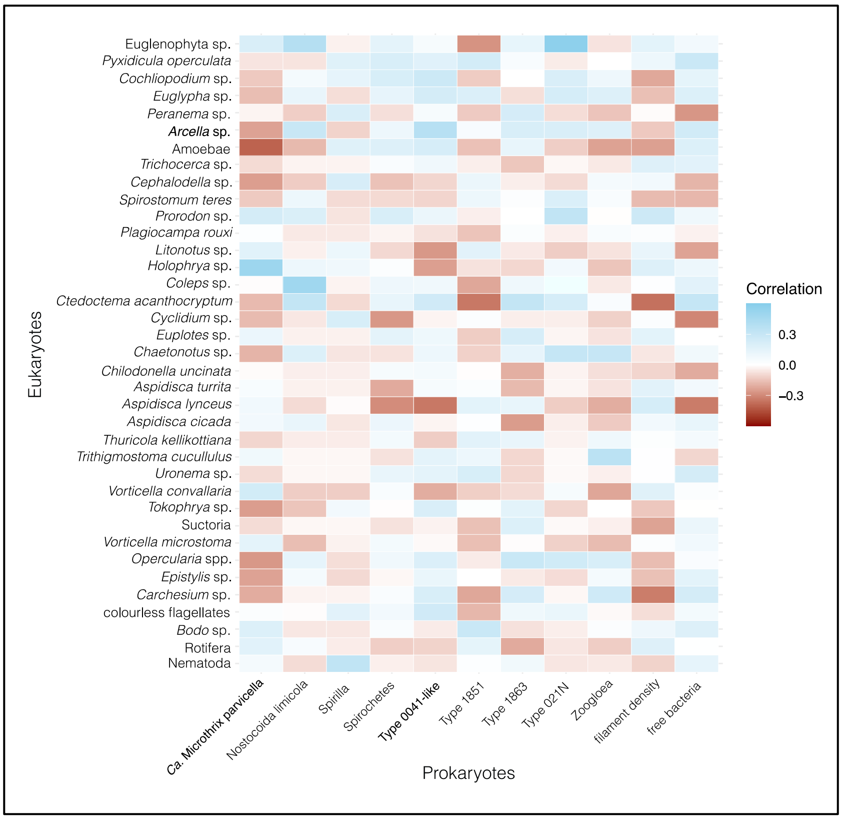


***Figure S4**: Heatmap depicting the results of a Spearman's rank correlation analysis for Dataset 2 between eukaryotic and prokaryotic taxa. Blue and red indicate positive and negative correlation coefficients, respectively. The strong negative correlations between Ca. M. parvicella and Arcella spp. as well as Amoebae suggesting an inverse relationship between the organisms.*

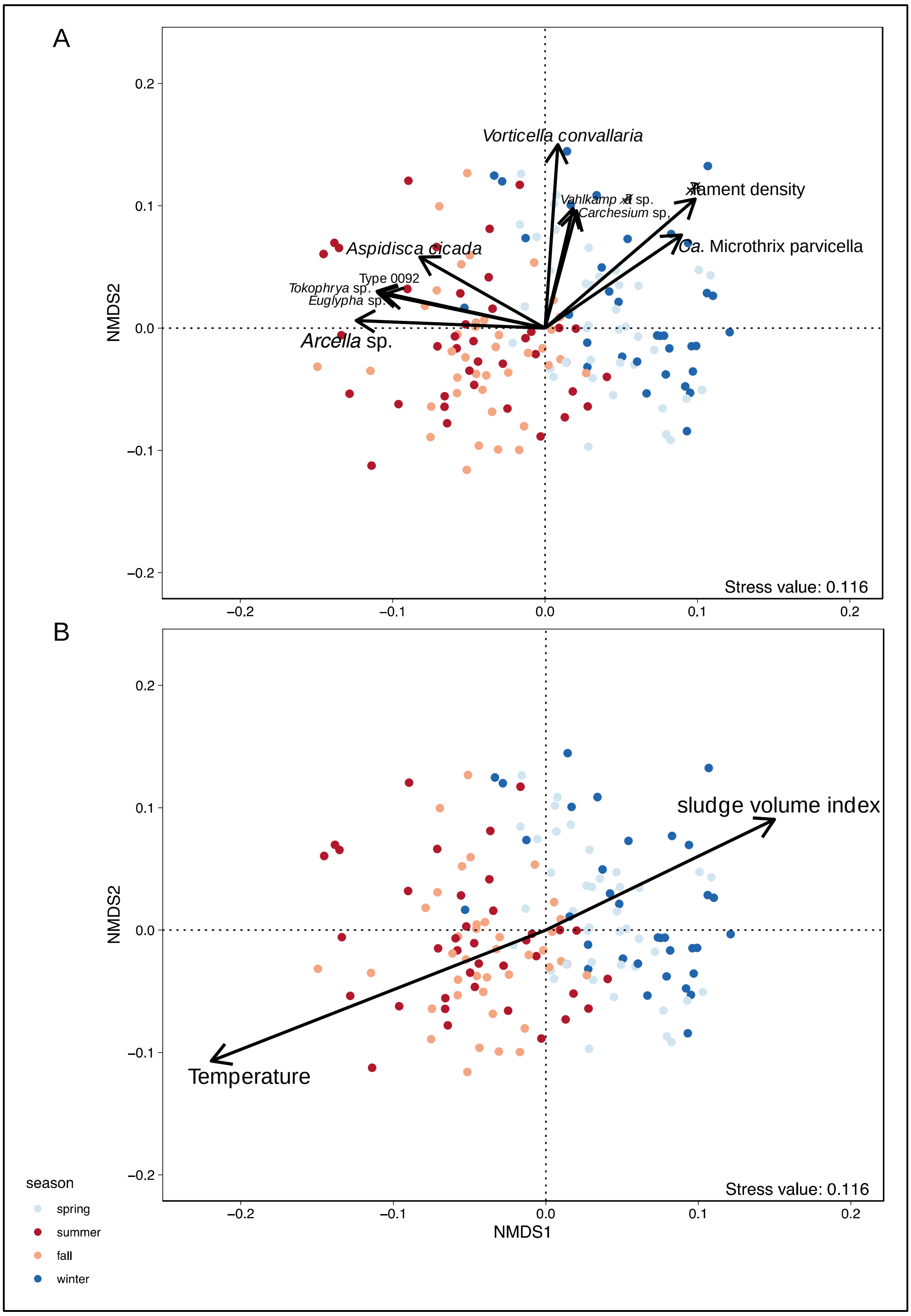


***Figure S5: Seasonal dynamics in the microbial communities of an aerated bioreactor (Dataset 3)***. *(A) Non-metric multidimensional scaling (NMDS) plot, with taxonomic organism variables fitted to the ordination space, illustrating the seasonal dynamics of the microbial community. The colours of the data points indicate the corresponding season (light blue = spring, blue = winter, red = summer, orange = fall). The length of each arrow represents the strength of the association between the corresponding organism and the ordination space. (B) Non-metric multidimensional scaling (NMDS) plot, with metadata variables fitted to the ordination space. Ca. M. parvicella and filament density shows a strong negative association with temperature and a positive association with the sludge volume index. Arcella spp show a more positive association with temperature.*

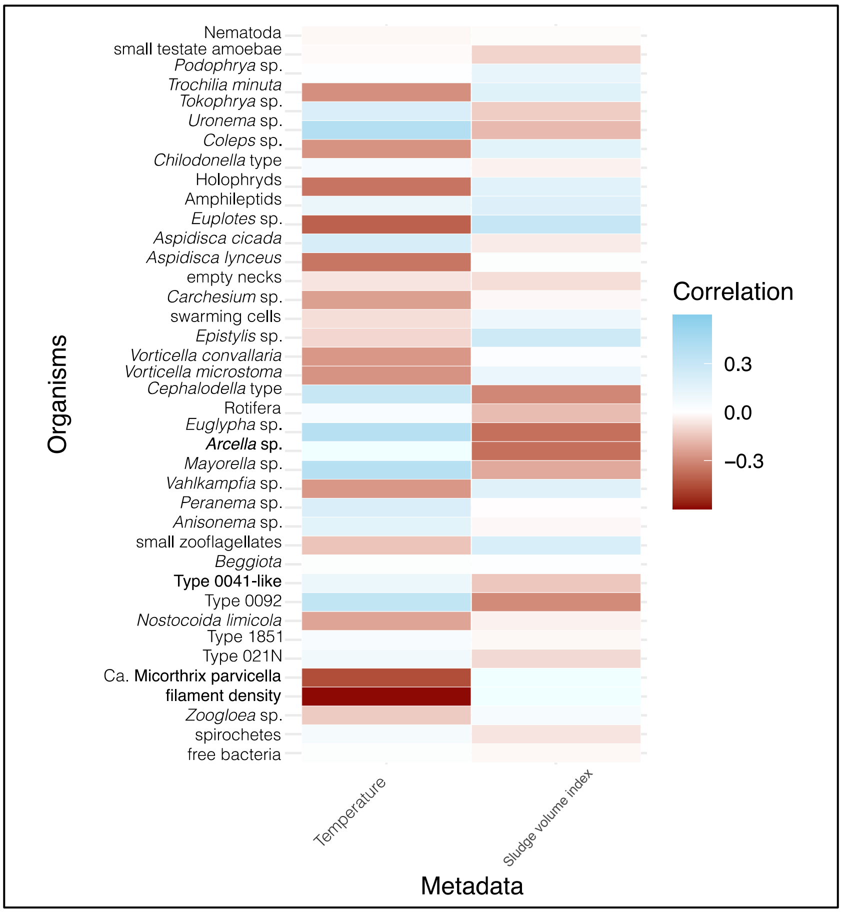


***Figure S6:*** *Heatmap depicting the results of a Spearman's rank correlation analysis for Dataset 3 between the observed taxa and associated environmental metadata. Blue and red indicate positive and negative correlation coefficients, respectively. Note the strong negative correlation between the abundance of Ca. M. parvicella, filament density and temperature, indicating a temperature dependence.*

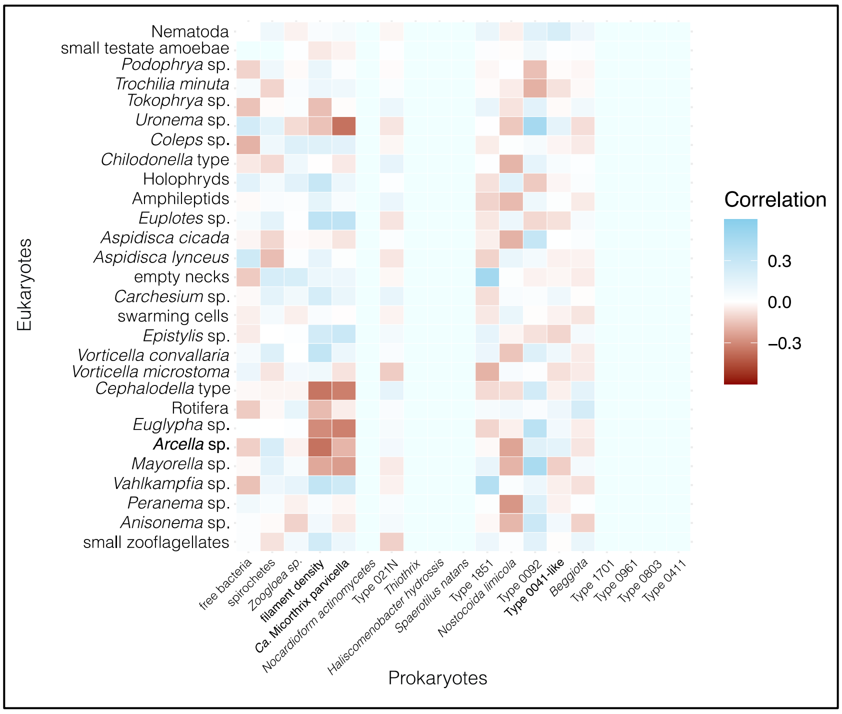


***Figure S7:*** *Heatmap depicting the results of a Spearman's rank correlation analysis for Dataset 3 between eukaryotic and prokaryotic taxa. Blue and red indicate positive and negative correlation coefficients, respectively. Note the strong negative correlations between Ca. M. parvicella, filament density and Arcella spp. as well as other amoebae suggesting an inverse relationship between the organisms.*

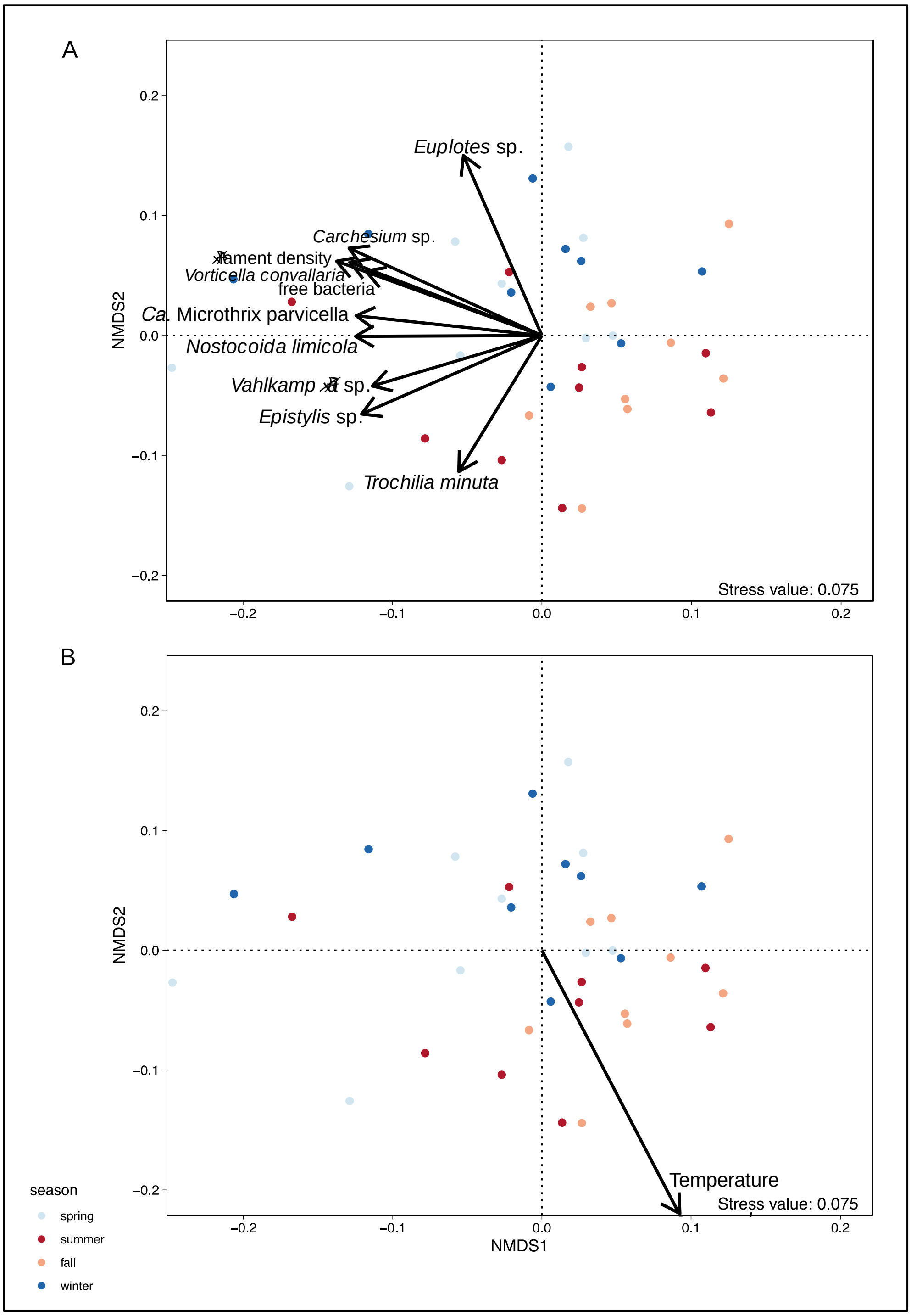


***Figure S8: Seasonal dynamics in the microbial communities of an aerated bioreactor (Dataset 4)***. *(A) Non-metric multidimensional scaling (NMDS) plot, with taxonomic organism variables fitted to the ordination space, illustrating the seasonal dynamics of the microbial community. The colours of the data points indicate the corresponding season (light blue = spring, blue = winter, red = summer, orange = fall). The length of each arrow represents the strength of the association between the corresponding organism and the ordination space. (B) Non-metric multidimensional scaling (NMDS) plot, with metadata variables fitted to the ordination space. Ca. M. parvicella shows a negative association with temperature.*

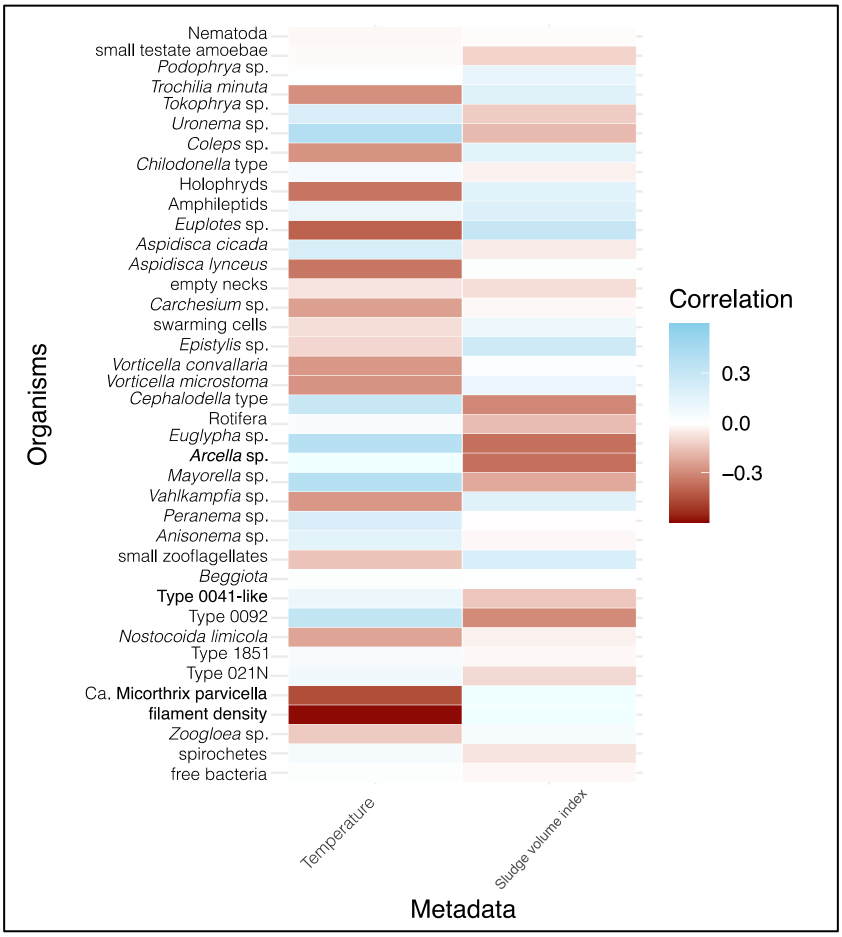


***Figure S9:*** *Heatmap depicting the results of a Spearman's rank correlation analysis for Dataset 4 between the observed taxa and associated environmental metadata. Blue and red indicate positive and negative correlation coefficients, respectively. Note the strong negative correlation between the abundance of Ca. M. parvicella, filament density and temperature, indicating a temperature dependence.*

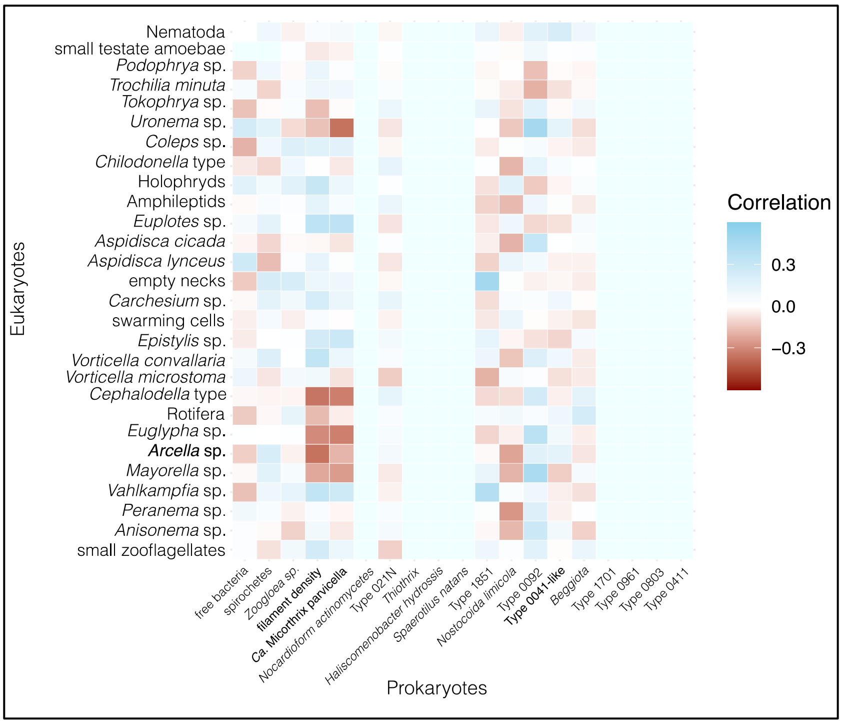


***Figure S10****: Heatmap depicting the results of a Spearman's rank correlation analysis for Dataset 4 between eukaryotic and prokaryotic taxa. Blue and red indicate positive and negative correlation coefficients, respectively. Note the strong negative correlations between Ca. M. parvicella, filament density and Arcella spp. as well as other amoebae suggesting an inverse relationship between the organisms.*

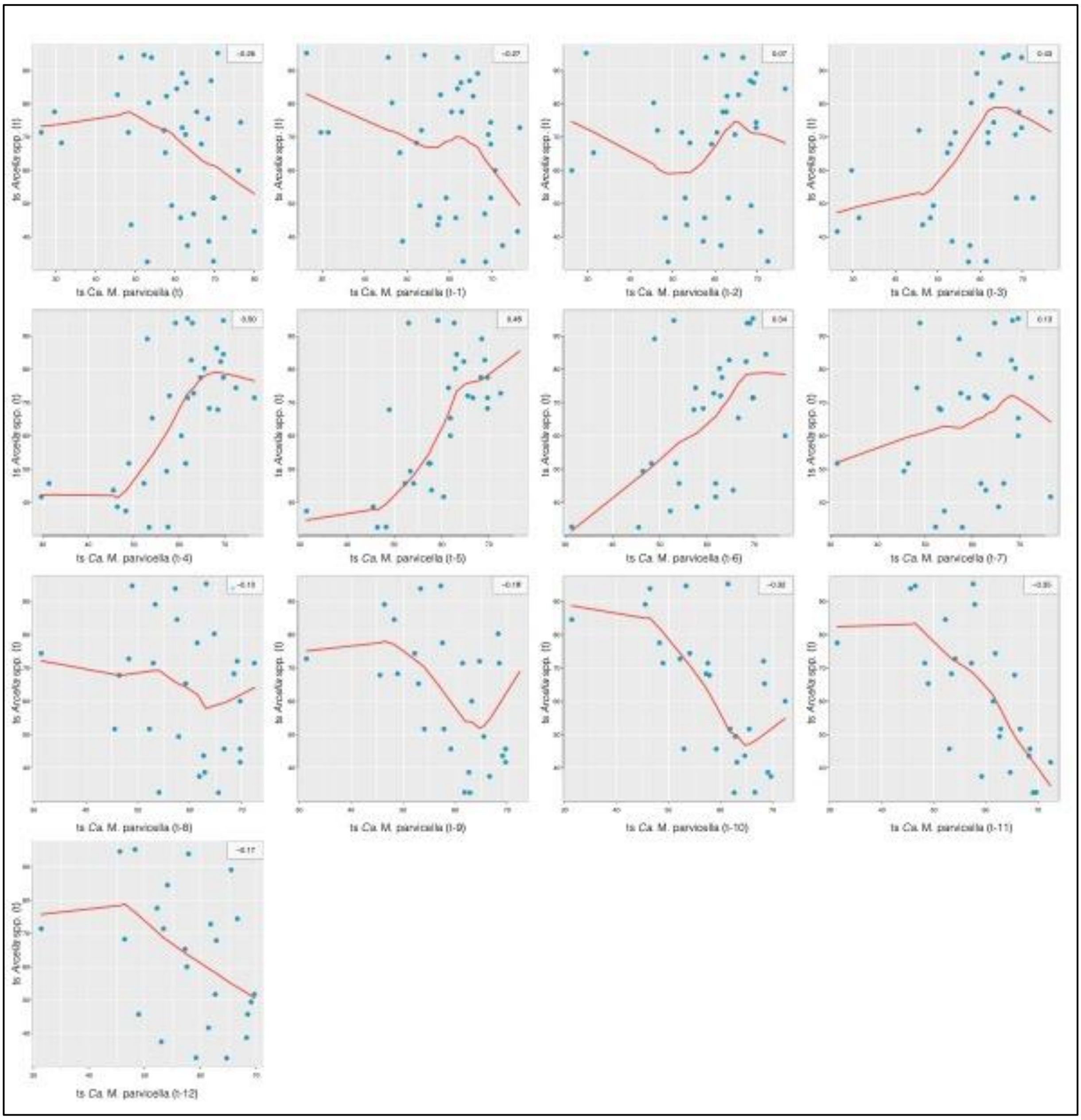


***Figure S11: Extended lag analysis of Ca. M. parvicella and Arcella spp..****This figure expands on Figure 5A, showing lagged correlations between Ca. M. parvicella abundance at previous time points (t-1 to t-12) and Arcella spp. abundance at the current time point (t). Each subplot represents a specific lag interval, illustrating how the temporal dynamics of Ca. M. parvicella influence the abundance of Arcella spp.. The analysis highlights the delayed predator-prey relationship consistent with Lotka-Volterra dynamics, with notable correlations observed at specific lag periods.*
